# Unveiling key links between behaviour and appearance in the evolution of camouflage

**DOI:** 10.64898/2026.05.07.722737

**Authors:** Yuri Fanchini Messas, George R. A. Hancock, João Vasconcellos-Neto, Martin Stevens

**Author notes:** Joint first authors (Yuri Fanchini Messas, George R. A. Hancock).

## Abstract

Behaviour is a key yet often overlooked component of animal camouflage and how it evolves alongside colour and morphology remains poorly understood. The repeated evolution of stick-like postures in spiders offers a useful framework for investigating the importance of behaviour for concealment, since matching the environment should rely on specific body forms and postures, not just colouration. We hypothesised that when spiders behaviourally align their body with the background orientation it should influence the shape, posture and colouration that best enhances camouflage. To test this, we used a genetic algorithm and human observers to evolve digital spiders to be harder to find. We evaluated how selection under three behavioural orientation treatments (aligned, random, and evolvable orientation) influenced spider capture time, background match (lightness and colour), posture, and body (cephalothorax and abdomen) dimensions. We found that spiders that behaviourally aligned with the background took substantially longer to find through evolving a better background match, and a more elongated posture and body shape than randomly orientated spiders. Our spiders mirrored the shape and posture adopted by numerous clades, illustrating how behavioural camouflage represents a key concealment strategy in structurally complex habitats, part of an interacting suite of traits that encompass successful concealment.

## INTRODUCTION

Animal coloration is a key trait that mediates interactions between organisms and their environments, influencing both concealment and communication in ways that significantly impact survival and reproduction (Caro et al. 2017; Cuthill et al. 2017). Camouflage has long provided striking examples of natural selection, yet many of its underlying mechanisms remain unexplored. While most research has focused on visual appearance, effective crypsis often relies on the integration of a variety of evolvable traits. Behaviour, for example, can be combined with colour and shape to facilitate successful camouflage by determining an organism’s location, orientation, and posture (Stevens et al. 2017; Stevens and Ruxton 2019). It is increasingly appreciated that successful background matching often relies on individual animals choosing background patches that better match the appearance of themselves or their young (e.g. Lovell et al. 2013; Camacho et al. 2020; Alothyqi et al. 2024; reviewed by Stevens and Ruxton 2019). However, the importance of behaviour in concealment in many taxa should go far beyond this. Classic studies on postural camouflage demonstrated that moths align their bodies with background features to enhance crypsis (Sargent 1965; Endler 1984; Grant and Howlett 1988; Webster et al. 2009; Kang et al. 2012; Webster 2015), cuttlefish dynamically adjust their arm positions to match different substrates (Barbosa et al. 2012), and even shorebirds modify posture to reduce detectability while balancing thermoregulation and predator avoidance (Timmis et al. 2022). Together, these and many other examples highlight that behaviour is integral to camouflage and may play an important role in shaping its effectiveness and evolution.

Structurally complex habitats, such as those with branched features, provide a valuable natural system for investigating camouflage mechanisms, as selection for crypsis in these environments has been observed to repeatedly lead to convergent evolution of similar appearances in unrelated species. Within such habitats, many animals rely on elongated or irregular body shapes, patterns, and postures to reduce detectability among twigs and branches. Species that rest or hunt on sticks – such as orthopterans, mantises, assassin bugs, owlflies, moths, and spiders – frequently display colours and body shapes that closely match those of their substrates (Edmunds 1976; Bian et al. 2016; Kelley et al. 2022). Most notable are thousands of species of stick insects (Phasmatodea) distributed worldwide, which have evolved both morphology and behaviour that resemble and sway like twigs (Robinson 1981; Bian et al. 2016). These examples illustrate a continuum of camouflage strategies ranging from less specialised background matching to masquerade and twig mimicry (Skelhorn et al. 2010a,b). However, while it is increasingly appreciated that behaviour and appearance work in sync to provide successful camouflage, how they do so, and how these traits evolve together, is poorly understood (Stevens and Ruxton 2019). Key uncertainties remain regarding whether behaviour or morphology evolves first, or both together in steps, and how important each is in preventing detection or recognition.

Spiders provide particularly suitable models for testing hypotheses about animal coloration, as they are highly diverse, abundant, and exhibit a broad range of colour patterns involved in several concealment strategies (Pekár 2014). Across continents and spider guilds, many lineages have independently evolved both appearance (e.g. elongated abdomen and colour matching the substrate) and resting postures (e.g. extended- or retracted-clustered legs and/or body alignment in relation to the substrate) that likely reduce detectability on sticks and twigs (Figure 1). These “stick-like” spiders are typically sit-and-wait predators distributed across at least 12 families (Araneidae, Archaeidae, Deinopidae, Desidae, Oxyopidae, Philodromidae, Pisauridae, Tetragnathidae, Theridiidae, Thomisidae, Senoculidae, and Uloboridae), which display diverse foraging strategies, from non-weavers to orb- and space web builders (see Supplementary Table S11). Zhang et al. (2015) showed using prey models that clustering the legs linearly, as is done by *Ariamnes cylindrogaster* Simon, 1889 (Theridiidae), reduces the rate of predation, potentially by masking the spider’s contour. Other species from at least seven families rest with their legs fully extended along sticks (Figure 1, Supplementary Material SI 5), suggesting that posture may act as a flexible behavioural trait that enhances concealment and potentially shapes body form under natural selection. However, the evolutionary function of this posture remains unexplored, and it is currently unclear for spiders, and many other taxa, how integral behaviour is to the evolution of specialised camouflage and its success.

**Figure 1.**
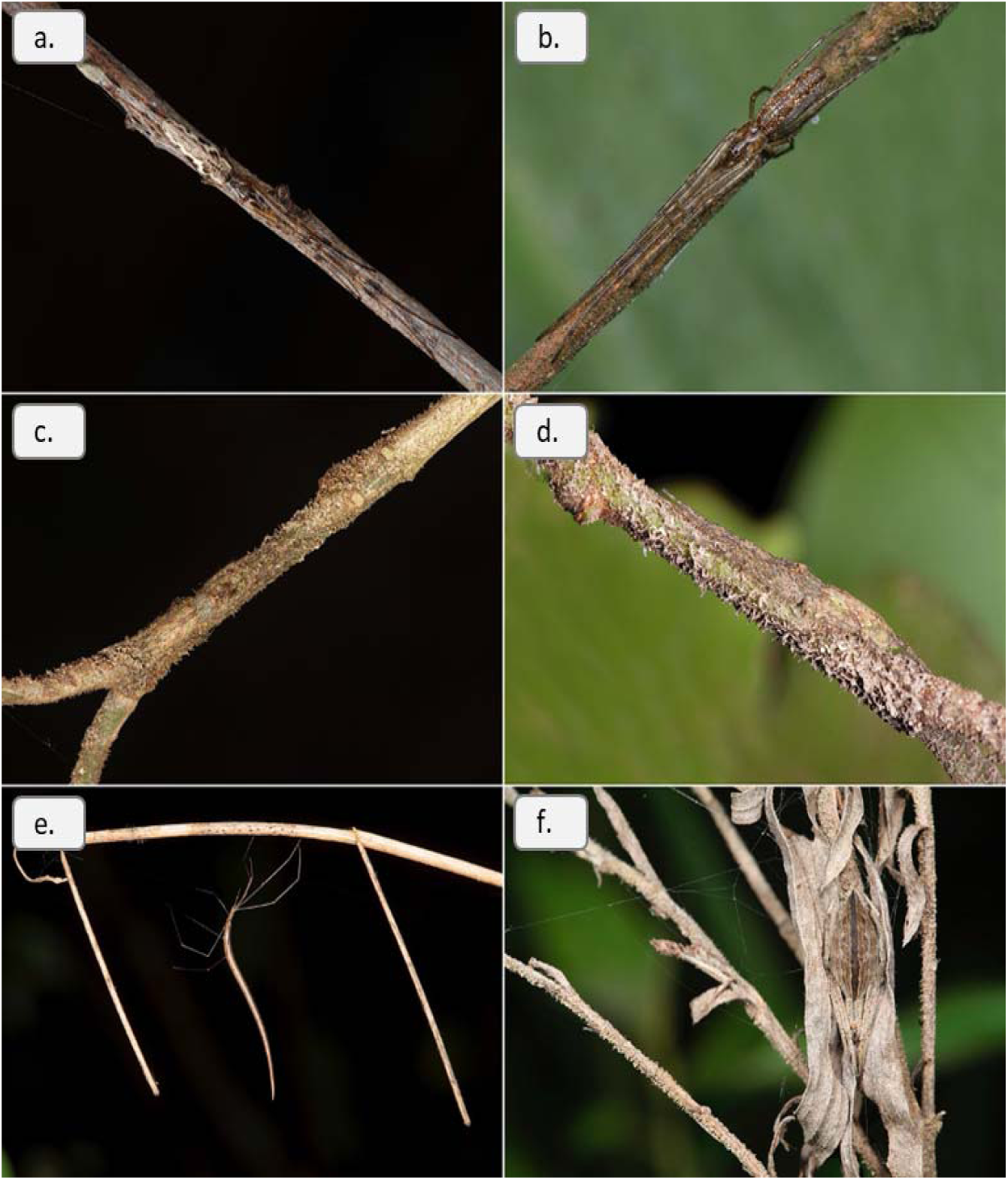
(a-d) Examples of stick-like spiders with elongated body shapes and their typical rest position, with the body aligned to the stick and legs extended linearly along the body axis, in branched environments: *Tetragnatha* spp. (Tetragnathidae) from (a) southeastern Brazil and (b) south United Kingdom; (c-d) dorsal and lateral view of a *Senoculus* sp. (Senoculidae) from southeastern Brazil. (e-f) Examples of different strategies of concealment of (e) the cobweb spider *Ariamnes* sp. (Theridiidae) sticking out perpendicularly to the branch and (f) the legs retracted and clustered close to the body of the orb-web spider *Eustala sagana* (Araneidae).

Since we cannot directly observe the evolution of the above traits, genetic algorithms and evolutionary simulations provide an excellent opportunity to understand the emergence and value of key camouflage traits (Mitchell 1996; Hamblin 2013; Hancock and Troscianko 2022). Genetic algorithms can be combined with citizen science experiments with digital animals to test hypotheses in evolutionary ecology (Bond and Kamil 2002; Bonney et al. 2014; Talas et al. 2017; Fennell et al. 2021; Briolat et al. 2024; Hancock et al. 2025). Thus far to our knowledge, no study has combined colour, morphology, and behaviour to test the adaptive significance of animal coloration, and studies of posture in camouflage are very rare, often just anecdotal descriptions or subjective observations. Here, we address this gap in our understanding of the role of behaviour in camouflage evolution by using digital imaging and computational modelling to design a digital evolution experiment to test how selection and behaviour shape appearance and posture in branched environments, using spiders as a model. Specifically, we tested (1) to what extent behaviourally matching the orientation of the background increases how hard our spiders are to detect, (2) whether behaviourally aligning with the background is required for stick-like postures to evolve, (3) whether aligning to the background relaxes selection on body size or shape, and (4), whether behaviour and its co-evolution with appearance improve the level of background match that evolves.

## MATERIALS AND METHODS

### Participants

We recruited 30 participants with normal or corrected-to-normal vision as volunteers from the University of Exeter. Before the trials, participants read a short instruction sheet and signed a consent form in line with the Declaration of Helsinki. The study was approved by the University of Exeter Bioscience Ethics Committee (application no. 2022/516616).

### Experiment design

We utilized the open-source CamoEvo toolbox (Hancock and Troscianko, 2022), implemented in ImageJ (Schneider et al., 2012), to simulate the evolution of spiders on branched backgrounds. We modified the existing animal camouflage pattern generator to be able to produce spiders with different postures (see Supplementary Material and Figure 2). Each population was programmed to evolve under one of three treatments, corresponding to three different behaviours: (1) random = the orientation of the spider was always random, (2) aligned = orientation always matched the branch the spider was on (orientations were pre-determined by experimenters), and (3) genetic = the orientation of the spider was determined as a decimal gene (0-1) encoding a linear range from 0-90 degrees. For each treatment we assigned 10 participants totalling 30 populations of spider. Our spiders were shown against photographs of branched habitats taken under controlled lighting conditions (see below) and each participant was tasked with finding the spiders as fast as possible. Each population consisted of 36 spiders, for each generation the fittest (highest capture time) 1/3^rd^ of spiders survived and reproduced in two rounds of recombination, resulting in a new population of 36 spiders. We expected our measure of fitness (capture time) to increase with successive generations (evolutionary time), and we measured camouflage from their behaviour (posture and orientation), morphology (length, width, and aspect ratio), and coloration (colour and lightness differences between spiders and their background).

**Figure 2.**
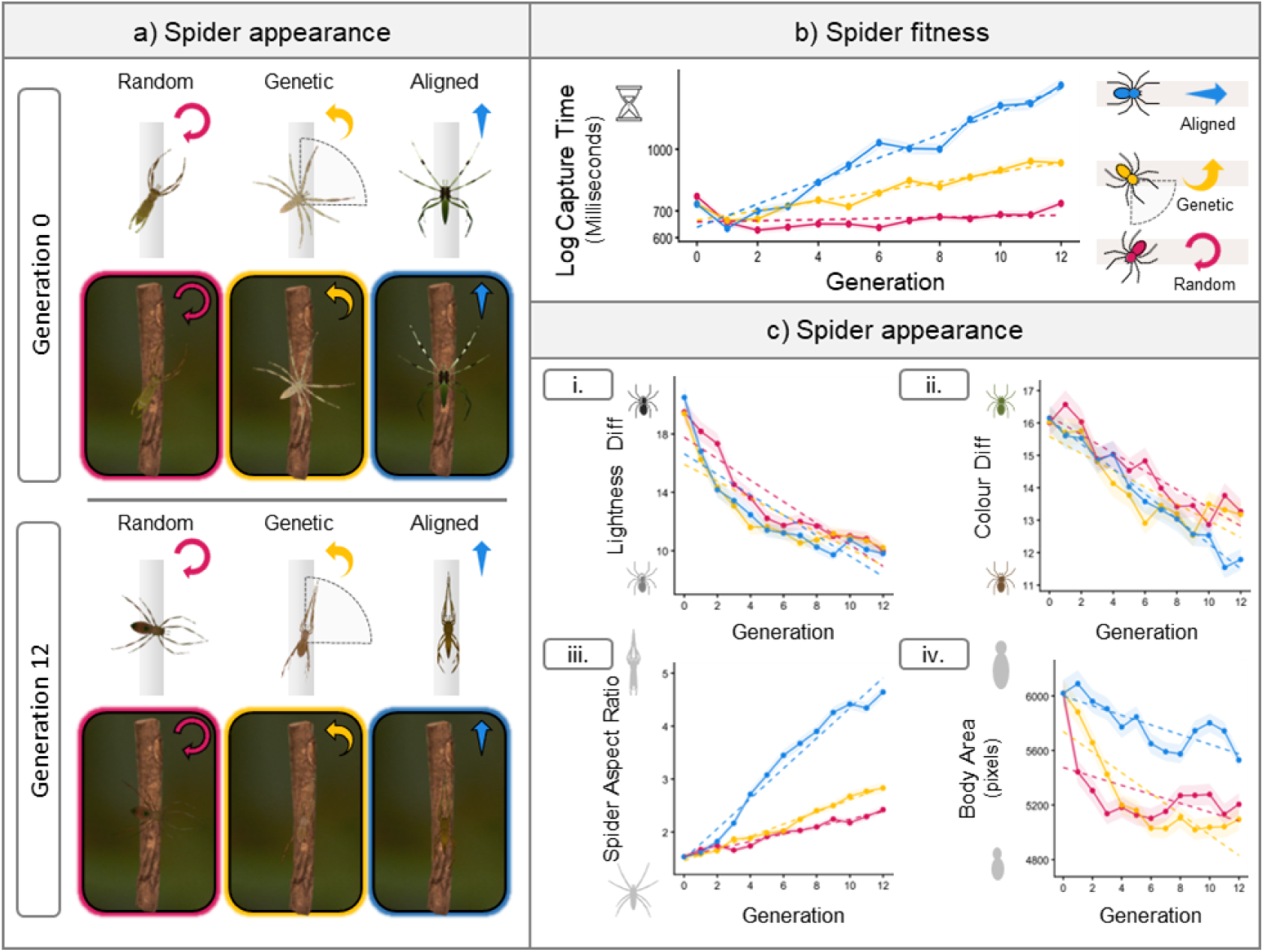
Spider targets evolved along 12 generations against branched backgrounds using ten different starting populations divided into (a) Example changes in colouration, posture and orientation for the behavioural treatments for the first and last generations for each spider behaviour. Each spider is shown both with and without a zoomed in section of the background. For a full set of examples, see the Supplementary Material section SI 4 (b) The change in the fitness measure (capture time) for each behaviour. (c) Quantification of changes in spider appearance, (c.i) the lightness difference (CIE L*) of the spider from the branch behind it, (c.ii) the colour difference (Euclidean CIE a* and b*) of the spider from the branch behind it, (c.iii) the aspect ratio of the whole spider, and (c.iv) the area of the spider’s body (cephalothorax and abdomen). For (c) and (d), dashed lines show the linear trend, while the solid lines show the change in mean value for each generation and the standard error (ribbon). Variation in starting values for (c.i) and (c.ii) were due to differences in background location.

### Background images and targets

To construct our game, we photographed natural branched habitats with an ASUS A002 smartphone and calibrated the images using 4.5 and 96.2% grey reflectance standards. Each experimental background image consisted of two merged layers, one of sticks forming a branched environment and the other of a vegetated background (see supplementary material for examples). We collected 120 random sticks in the woodlands of Falmouth (Cornwall, UK), where stick-like spiders, such as the long-jawed orb weavers (e.g. *Tetragnatha*, Figure 1b) occur, and subsequently we randomized their orientation, number, and type on a grey background. We positioned the smartphone on a tripod with the lens perpendicular to the sticks inside a Neewer diffuser tent of 1.5 x 1.5 m to obtain 36 different photographs. We also photographed 36 random vegetation backgrounds where spiders could occur. We calibrated the colours and rescaled all images using the micaToolbox package (Troscianko and Stevens 2015) within ImageJ. Then, we removed the background from the stick images, re=cropped them to 1800 px × 2400 px and merged the two layers (stick and background) using Adobe Photoshop (version 23.5.0, Adobe™).

For each background, we used CamoEvo 2.0’s ‘spawn map’ feature to label where on the background images spiders were allowed to occur. The ‘perceived’ angle of the sticks was also labelled by marking lines along the stick and measuring the angle in ImageJ (see Supplementary Material SI 2). This was used to ensure spiders would not appear / ‘spawn’ in areas that were not in contact with the sticks and allowed the aligned and genetic treatments to orient relative to the background. The game stimuli consisted of spiders made by CamoEvo (max 270 px, downscaled from a 400 px image) shown against random backgrounds from a pool of 36 substrate images (2400 px, 1800 px). Targets were allowed to occupy a CIELAB space (CIE 2007; Renoult et al. 2017) with a lightness range of 0 to 100, A (green-red) range of −25 to 35, and B (blue-yellow) range of 5 to 50.

### Evolution experiment

Using CamoEvo, we created 10 unique starting populations of 36 spiders matched across our three behavioural treatments. This was to help ensure that differences in evolved lines were due to selection as opposed to random differences in the starting phenotypes. Each participant was tasked with evolving their assigned population and behavioural treatment for 12 generations. The number of generations was determined based on previous uses of CamoEvo, to keep the duration of the experiment under an hour and as the goal was to test initial differences in selection as opposed to global optima for phenotype. For each generation, CamoEvo automatically paired each spider with one of the 36 background images (1 of each background per generation) to create a total of 36 stimuli, totalling 468 trials per participant.

Each participant was given one hour to complete the experiment in a room illuminated only by the monitor, located in the Centre for Ecology and Evolution at the University of Exeter. Using the same computer allowed us to standardise the computer display (ProLite e2283hs 21.5” monitor), operating system (Microsoft Windows), capture method (mouse) and viewing distance (50 cm). Before each trial started, the volunteers were instructed by CamoEvo to position their cursor at the centre of the screen and to only move it after seeing the target, which they then had to click on spiders as quickly as possible within 15 seconds per trial. After each generation was completed, a new population was automatically generated. Ranked capture time was used as the measure of fitness. Previous work has shown that the first target typically takes far longer to find than subsequent ones (Troscianko et al. 2018). To help control for this novelty effect, participants were shown a sample slide on a printed instruction sheet prior to each game.

### Measures

For each trial stimulus we measured the mean CIE L* (lightness), CIE a* (green-red), and CIE b* (blue-yellow) colour channels for the spider and the background immediately behind the centre of the spider (10 px radius circle), as well as the local surround (circle 2x area of spider). These were used to calculate the difference in mean lightness and in colour (Euclidean a* and b*) from the background. The expectation being that lower difference values should equate to better camouflage, as has previously been shown(Nokelainen et al. 2019; Hancock and Troscianko 2022; Hancock et al. 2025). We also measured the shape (height, width, and aspect ratio) of the spider as an estimate of posture and of the spider’s body (cephalothorax and abdomen, without the legs). Narrower twig-like postures increase the aspect ratio of the spider (see Supplementary Material SI 1).

### Statistics

All statistical analyses were conducted in R version 4.2.2, using the package lme4 (version 1.1-27.1) to fit linear mixed models (Bates et al. 2015; R Core Team 2023). To test whether our behavioural treatments influenced camouflage, we used log(capture time), lightness difference, and colour difference as our response variables, and the behaviour assigned to the population and its interaction with the generation number as our predictor variables. The expectation being that if camouflage improves, then capture time should increase and both lightness and colour difference should decrease across generations. The starting population and the background against which the target spiders were shown were used as random effects. To compare differences between behaviours, we used the emmeans package to conduct Tukey post hoc comparisons (Lenth et al. 2019). An example lme4 coded model is given below:

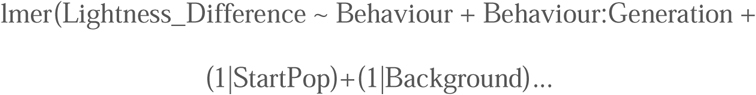

Likewise, to investigate the influence of behaviour on posture and body shape, we used the aspect ratio of the whole spider and the bounding area (width * height) of the spider’s body as our predictor variables. Lastly, to determine whether the orientation match (lower angular difference) co-evolved with better camouflage, we first tested whether our genetic treatment evolved a better angle match to the background than the random using our behaviour treatment (random or genetic), generation, and their interaction as our predictors and angular difference as our response variable. Then we tested whether spiders with a better angle match and narrower postures had a higher capture time, and thus better camouflage, for the genetic treatment compared to the random, by using behaviour, angle difference and spider aspect ratio as our predictors.

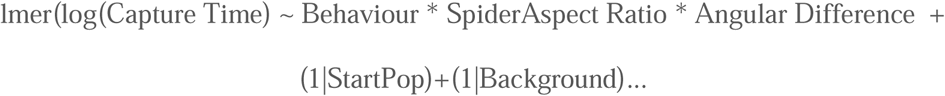

These models included only our random and genetic treatments, because the aligned treatment always matched the background, resulting in no variation in angular difference. As capture time, aspect ratio, and angular difference were on different scales, these variables were rescaled to a mean of zero and a standard deviation of 1, for the last model. Model assumptions were evaluated for all models by inspecting residuals for normality and homoscedasticity.

## RESULTS

For the full model outputs and Tukey post hoc comparison tables, please see our Supplementary Material section SI 3.

### Effects of behaviour on camouflage

For our three behaviour treatments (Figure 2), capture time was found to only significantly increase for spiders evolved under our background-aligned and genetically controlled orientation treatments (Random:Generation, N-Gen, β = 0.003, t_13990_ = 1.64, p = 0.101; Genetic:Generation, N-Gen, β = 0.028, t_13990_ = 14.14, p = 0.101; Aligned:Generation, N-Gen, β = 0.067, t_13990_ = 33.98, p = 0.101), with our aligned treatment evolving the greatest capture time (Post Hoc | Genetic vs Aligned, z = -18.62, p < 0.001). The initial mean capture time (gen = 0) was 848 milliseconds, whilst in the final generation (gen = 12) the means were, random = 851, genetic = 1111, and aligned = 2054 milliseconds.

Despite our use of examples to reduce the influence of learning, capture time was observed to initially decrease between generation 0 and 1 and then increase again at varying rates, suggesting that volunteer learning initially outpaced camouflage evolution. This observation is supported by the fact that the colour and lightness difference from the background still significantly declined for the random treatment (Random:Generation, N-Gen, β = -0.700, t_13990_ = -17.51, p = 0.101) (Figure 2d), though to a significantly greater degree for the aligned and genetic treatments for lightness (Post Hoc | Random vs Genetic; z = 4.30, p < 0 .001; Random vs Aligned, z = 4.35, p < 0.001; Genetic vs Aligned, z = 0.05, p = 0.999) and colour (Post Hoc | Random vs Genetic; z = 3.25, p < 0.001; Random vs Aligned, z = 4.66, p < 0.001; Genetic vs Aligned, z = 1.40, p = 0.339). No significant difference was observed between the genetic and aligned treatment for either lightness or colour difference. Additional analyses of camouflage measures found that for the random treatment, compared to aligned, lightness difference from the local surround (area 2x the size of the spider) was a better predictor (lower AIC) of capture time than the area behind (see Supplementary Material SI 3). But for colour, only the immediate background had any predictive value with local colour match having no significant effect on capture time if the orientation was random, and local colour difference significantly increased rather than decreased capture time if the spiders were always aligned.

### Effects of behaviour on posture and body shape

The aspect ratio of the spiders, used as an indicator for how stick-like the posture of the spiders was (see Supplementary Material SI 1), significantly increased for all our genetic and aligned treatments, with the lowest increase for the random and the greatest increase of the aligned (Random:Generation, N-Gen, β = 0.07 t_13990_ = 12.66, p < 0.001; Genetic:Generation, N-Gen, β = 0.12, t_13990_= 20.83, p < 0.001; Aligned:Generation, N-Gen, β = 0.12, t_13990_ = 51.45, p < 0.001) (Figure 2diii). Spiders evolved under always aligned orientations, developed a higher aspect ratio than those where orientation was genetic (Post Hoc Genetic vs Aligned, z = -15.02, p < 0.001). Spiders evolved with random and genetic orientations were rapidly selected to have smaller body areas, while the aligned treatment declined in area, but to a considerably lesser degree (Post Hoc | Random vs Genetic, z = -0.195, p = 0.9792; Genetic vs Aligned, z = -16.399, p < 0.001; Genetic vs Aligned, z = -16.203, p < 0.001). This decline in area was primarily due to a drop in body width, with no significant difference observed between generations for body length in aligned treatment populations (see Supplementary Material SI 3).

### Effects of angular difference on fitness

The angular difference between the spider and the background for the genetic treatment significantly decreased across generations, with spiders becoming more aligned with the background (Genetic:Generation, N-Gen, β = -2.58, t_9347_ = -27.92, p < 0.001), unlike the random treatment, which did not improve (Random:Generation, N-Gen, β = 0.14, t_9347_ = 1.52, p = 0.128) as it was not an evolvable trait (Figure 3). Orientations that had a lower angular difference from the background (closer to zero) (Scaled AngleDifference, β = -0.12, t_9312_ = - 8.57, p < 0.001) and with narrower postures (Scaled SpiderAspectRatio, β = 0.20, t_9323_ = 12.44, p < 0.001) had higher fitness for both random and genetic behavioural treatments. The Pg.14 slope of fitness for angle difference was also steeper when the posture was narrower (Scaled SpiderAspectRatio:AngleDifference, β = -0.12, t_9323_ = -7.96, p < 0.001) and when the spider’s orientation was genetic (AngleDifference:Genetic, β = -0.04, t_9319_ = -3.56 p = 0.03). Remarkably, fitness for the genetic treatments was still higher than the random, regardless of interaction with angle and posture (Genetic vs Random, β = 0.129 t_9309_ = 9.31, p < 0.001).

**Figure 3.**
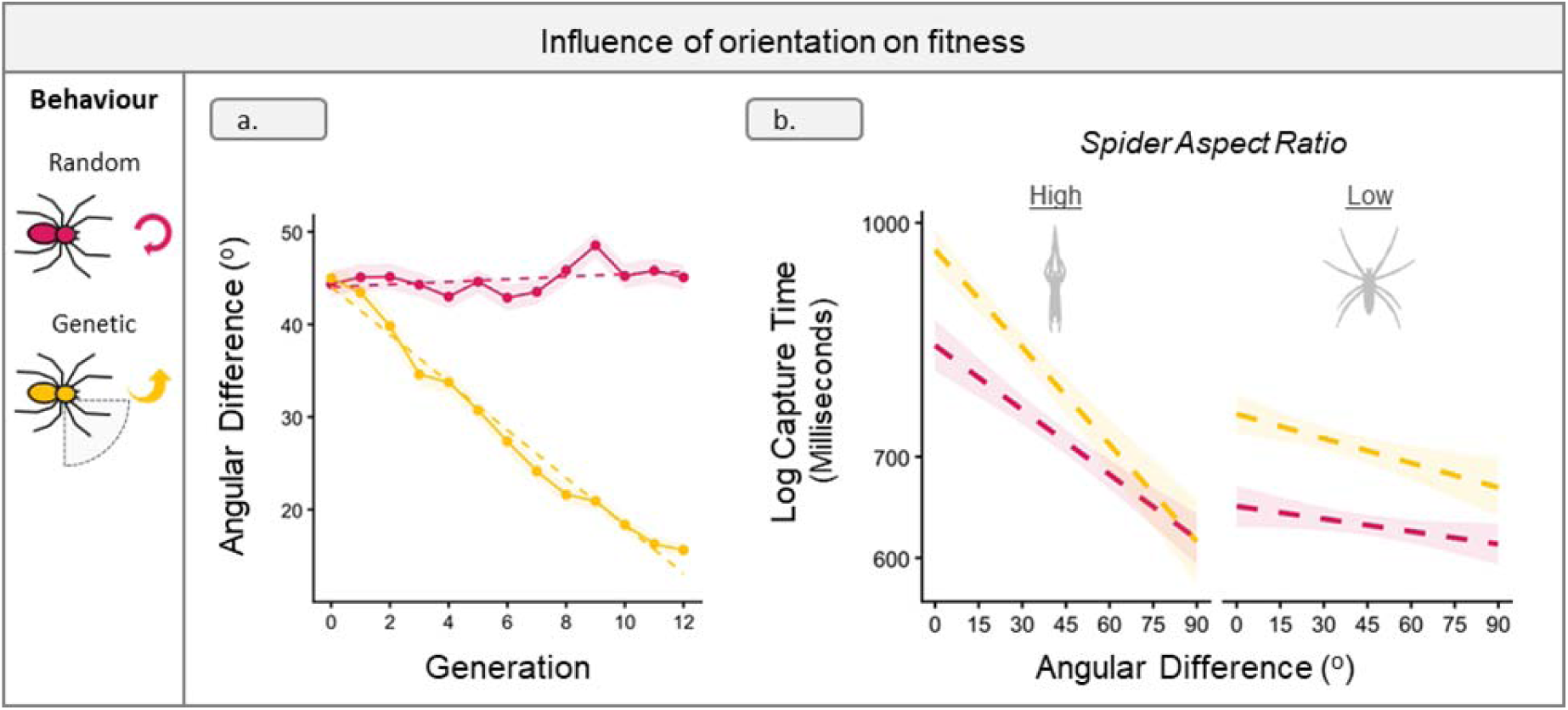
(a) Change in angular difference across generations for the two behaviours that had variable angles (Random and Genetic). (b) Influence of angular difference, and posture on log capture time for the different behaviours. Posture is split by high (> median) and low (<= median) spider aspect ratio. For (a) and (b), dashed lines show the linear trend, while the solid lines for (a) show the change in mean value for each generation and the standard error (ribbon).

## DISCUSSION

Here, we demonstrate, by using CamoEvo to digitally evolve camouflage with genetic algorithms (Hancock and Troscianko 2022), that behaviour plays a key role in facilitating spider camouflage in branched habitats. The capture time (fitness) of our digital spiders was increased by the ability to behaviourally align with the orientation of the branch the spider was on, and especially when spiders possessed postures that stretched the legs longitudinally, such that they did not splay outwards beyond the branch. Even when allowed to evolve alternative orientations, selection favoured orientations that matched the background structure, suggesting that, at least within the phenotypic space of our spiders, this was the best strategy for the backgrounds provided. In the absence of the ability to behaviourally align with the background, camouflage was found to be less effective, since spiders with a greater angle difference from the background were not only easier to find, but also were unable to evolve as effective a level of background match within the number of generations allocated. This supports the hypothesis that spiders camouflage against branched habitats by locally clustering with their perch (Zhang et al. 2015).

When spiders couldn’t behaviourally match orientation, the stick-like elongated posture, observed for at least 12 families of real spiders (see Supplementary Material SI 5), did not as consistently evolve, though their posture still became narrower (see Figure 2). Meanwhile, if the spiders always aligned to the background or evolve a matching orientation, then a more stick-like posture typically evolved. The rate at which stick-like postures increased in abundance within the populations was considerably faster if the spiders matched the orientation of their branch by default. This implies that behavioural control of orientation is a necessary precursor to evolving stick like postures, and must either exist first or co-occur with posture to be effective. Further to this, camouflage coloration functioned more effectively and evolved a more specialised match to the sticks as opposed to the surround when behavioural alignment could occur. Within habitats with regions that are highly directional, e.g. branched habitats, or are locally clustered in colour, behaviour is likely integral to the development of specialised and highly effective camouflage, as opposed to more general background matching and salience reduction (Hughes et al. 2019; Briolat et al. 2021; Murali et al. 2021).

Previous research using prey models has shown that leg clustering rather than body size predicts survival (Zhang et al. 2015), yet our work found a large body size to be strongly selected against when the spider’s body was allowed to orient away from the angle of the branched substrate the spider was on. This was true for both spider populations that evolved with random orientations and with angular differences from the background encoded by a gene. This was likely due to the abdomen sticking out from the branch when oriented away from it. However, for digital spider populations that always aligned with the background, selection against a larger body area was greatly reduced. Subsequent analysis showed that the width of the body declined to a lesser degree and the length did not significantly decrease (see Supplementary Material SI 3). Branch-dwelling spiders across multiple taxa have been found to have thin but long bodies. This alteration in body shape may reflect the need to maintain a large abdominal volume for other factors that influence fitness, such as gravidity (reproductive output) (Head 1995), whilst maintaining crypsis by reducing the visible area. While other non-exclusive factors, such as thermoregulation also select for elongation of the abdomen as it reduces sun exposure, as shown by (Ferreira-Sousa et al. 2021) for orb-web spiders. By using behaviour, spiders may mitigate the survival cost to crypsis of having a larger body and co-evolve shapes that support their behaviour. Variation in perch size and width may similarly constrain body size and shape for other taxa, such as lizards, due to crypsis; independent of locomotory and alternative ecological constraints (Steffen 2009; Jones and Jayne 2012; Corral-López et al. 2017).

Increasingly research has shown that a simple change in orientation can have a dramatic effect on detectability. Whether it is moths aligning their patterning to the background or animals adjusting their orientation with respect to the source of light to reduce shadows (Kang et al. 2012; Mavrovouna et al. 2021). Our work builds upon this by suggesting that previous assessments of camouflage effectiveness may have been influenced by overlooking behaviours that shape the success of different camouflage phenotypes. Amongst spiders, numerous bark-dwelling spiders (e.g. Selenopidae, Hersiliidae and Ctenidae) exhibit medial body stripes together with disruptive barring on legs which are often held in a splayed posture. Striped patterns have been shown to vary in effectiveness depending on the background structure and or lighting (Tsurui et al. 2010; Hancock et al. 2025). For animals with directional patterns and/or body shapes, the relative orientation between an organism and its background can be a key determinant of its detectability and survival. Failing to take orientation and behaviour into account is likely to influence the relative differences in camouflage performance between different animal patterns used in past digital camouflage experiments. For instance, striped patterns may become more generalisable or effective if the animal is allowed to adjust its orientation and position within its background (Kang et al. 2012; Briolat et al. 2021). Further investigations are also required to determine how different species alter their position, orientation, and posture when attempting to hide in different environments (Barbosa et al. 2012; Timmis et al. 2022).

While our study focused primarily on posture and shape, similar methods can be used to more intently explore possible influences of behaviour on camouflage coloration; for example, exploring the co-evolution of orientation and pattern shape in moths which have a more fixed appearance and less postural variation (Kang et al. 2012). Here, we would predict that allowing a digital moth to modify its orientation to the background would influence the directionality (stripiness) and orientation of its pattern that evolve. Background structure should also influence optimal behaviour, such that backgrounds with less pronounced directional features may not favour behavioural adjustment of orientations. Animals that exist in habitats with clustered appearances, such as patches of different vegetation and substrate, frequently possess intraspecific variation in appearances and/or colour-pattern types that they can change between, for example chameleon prawns and numerous cephalopods (Shohet et al. 2007; Barbosa et al. 2008; Green et al. 2019). Future work should consider exploring the influence of behaviour and background choice on the evolution and disparity between colour-pattern variants and their substrate, including the structure of the local environment.

A limitation of our study was the use of 2D digital as opposed to 3D environments, and the resulting extent to which the splayed postures of the spider jutted out beyond the line of the background, as we did not stimulate them gripping the sticks. For real spiders, the tarsus of the legs would typically be in direct contact with their substrate, for balance and locomotion, except when moving or displaying. This may have exacerbated the cost to crypsis of having a more splayed posture, though in turn this effect could have been nullified by the ability to see the curve of the limbs jutting out above the spider and/or branch when viewed from lower angles relative to the spider’s dorsum. Follow-up experiments, using 3D spiders, either within a virtual-reality environment (Augustyn et al. 2008; Matchette et al. 2018) or conducted in the field (Zhang et al. 2015; Briolat et al. 2024), would help to further validate our results in the context of spider orientation and posture. The density of branches, their width, and the variability of angles may also influence which posture is most effective and the degree to which orientation matching is selected for.

Our results showcase the remarkable relationship between behaviour, body shape, and animal coloration. This has broad implications not just for spiders but for animals that are capable of behaviourally altering their orientation and body shape/posture. Numerous invertebrates (e.g. spiders, mantids, moths, stick insects) and vertebrates (fish, reptiles, birds, mammals) have been observed to alter their postures or orientation when hiding against different substrates (Robinson 1981; Barbosa et al. 2012; Timmis et al. 2022). Without this behaviour, the effectiveness of their camouflage for concealing them from predators is likely to be greatly reduced. Our work here also highlights that evolution may impose strongest selection on certain traits first (e.g. posture or orientation) and then refine other features thereafter (e.g. colour), rather than necessarily acting on all traits similarly from the outset. By using digital evolution to evaluate optimal solutions and to heuristically search across wider phenotypic spaces, we can start to explore and unravel the intricacies of what drives the variability we see in the camouflage of animals. Subtle differences in behaviour may result in profoundly different camouflage strategies. Meanwhile, by allowing behaviour itself to evolve, we may better understand how animals can situationally enhance their camouflage or generalise across environments. Similarly, the co-evolution of shape and colour for masquerade, and other forms of mimicry (Jones et al. 2013; Kelly et al. 2021), may also be explored in this manner.

## CONCLUSIONS

Behaviour is increasingly appreciated as a critical component of successful concealment, spanning examples from where birds choose to lay their eggs, how moths align on trees, to the way some insects sway with vegetation movement (Kang et al. 2012; Bian et al. 2016; Stevens et al. 2017). One of the major recent gaps in understanding has been in the relative importance of behaviour and morphology in successful camouflage, and how these varied aspects of the overall defence evolve over time. Our work here illustrates that both behaviour and morphology are critical, but that the value of each is context dependent in line with the environmental features and the animal’s current biology. Overall, the ability of animals to adjust their orientation and posture can make a striking difference in the level of camouflage afforded by their morphology and in turn shape evolved appearance. This is demonstrated by our digital simulations and supported by the resemblance of our evolved phenotypes to the behaviours and morphologies exhibited by a variety of spider families that occupy habitats with a similar structure.

## DATA AVAILABILITY

The CamoEvo plugins, dataframes and R code supporting the findings of this study are available on Figshare: https://figshare.com/s/0e53af466dc62ee83b93

## AUTHOR CONTRIBUTIONS

Y.F.M. and G.R.A.H. contributed to the analysis and preparation of figures. G.R.A.H. created the custom ImageJ + CamEvo scripts used to generate the digital evolvable spiders and to measure their appearance. Y.F.M. collected and calibrated the images used to generate the backgrounds and recruited the volunteers for the experiment. All authors contributed to writing, editing, and reviewing the manuscript. Y.F.M & G.R.A.H. designed the study and experiments with input from MS.

## ACKNOWLEDGMENTS

We are extremely grateful to all the volunteers who participated in the experiment.

