## Supplementary Material for "Unveiling key links between behaviour and appearance in the evolution of camouflage"

### SI 1: Spider Generation

#### SI 1.0: Intro

A unique phenotype generator derived from the pre-release of the [CamoEvo V2.0](https://github.com/GeorgeHancock471/CamoEvo-v2.0-2022_Plugins) code was used for the spiders in the SpiderEvo game. This generator produced not only the patterns of the spiders but also the mask used to give them their shape and posture (Fig. S1).

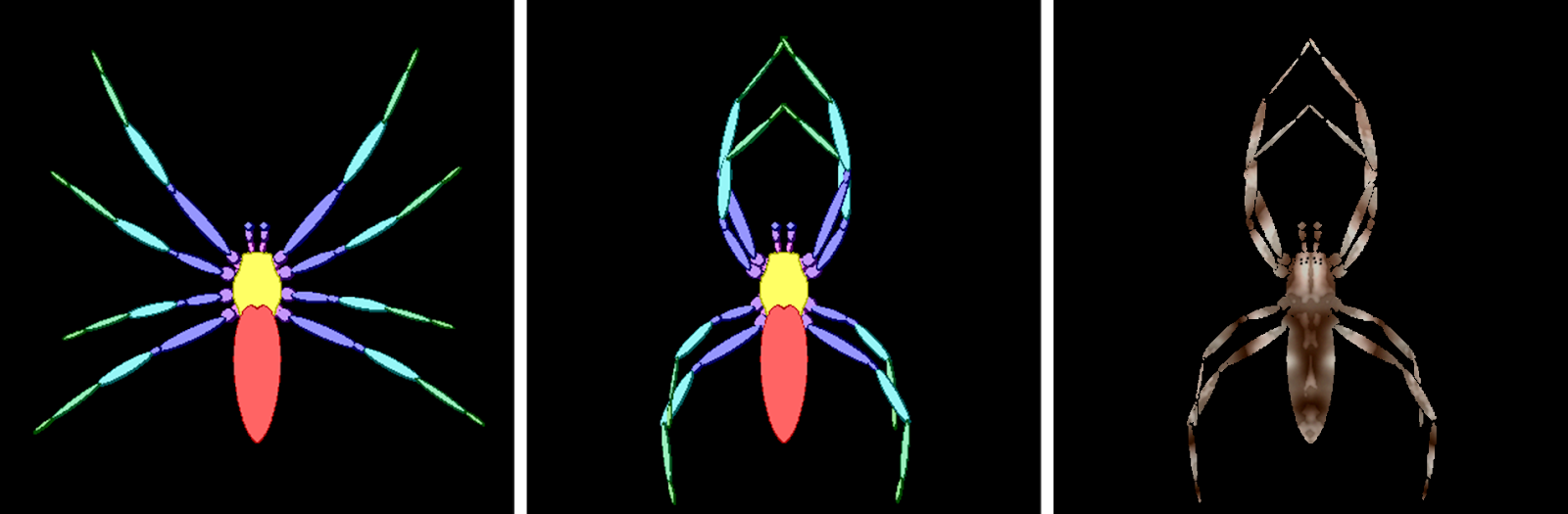

**Fig. S1.** Genetically controlled shape (left), posture (centre) and patterns (right).

The SpiderEvo masks were generated by editing the abdomen’s shape, the body’s size, and the legs’ width and angles (Fig. S2). Each spider started as a base cephalothorax shape. The abdomen was constructed using an oval, the width and height of which were adjusted genetically. By angling and mirroring the oval, pointed and round abdomens could be made. The cephalothorax and abdomen could be scaled up or down together. The length of the legs was constant to ensure that all spiders could have the same bounding area (posture determines bounds), and the width (thickness) of the legs was controlled independently from the body. See Fig. S3 for example scales.

The legs of the spiders were made by using a base-splayed limb configuration. Each limb consisted of 7 segments, barring the palps, which only consisted of three. The same gene influenced both leg and palp thickness. The posture of the legs was altered by using three genes: angle, strength and bend (Fig. S2). Angle determined the intended orientation; strength changed the ratio between the intended and the original splayed orientation, and bend caused the legs to curve towards the centre. The legs were split into two groups: group 1, the front p1 and p2, and group 2, the hind p3 and p4. Members of the same group shared the same posture genes. This system limited the number of genes needed and helped prevent impossible leg configurations but reduced the number of possible leg configurations. See Fig. S4 for example postures.

#### SI 1.1: Spider Shape & Posture Generation

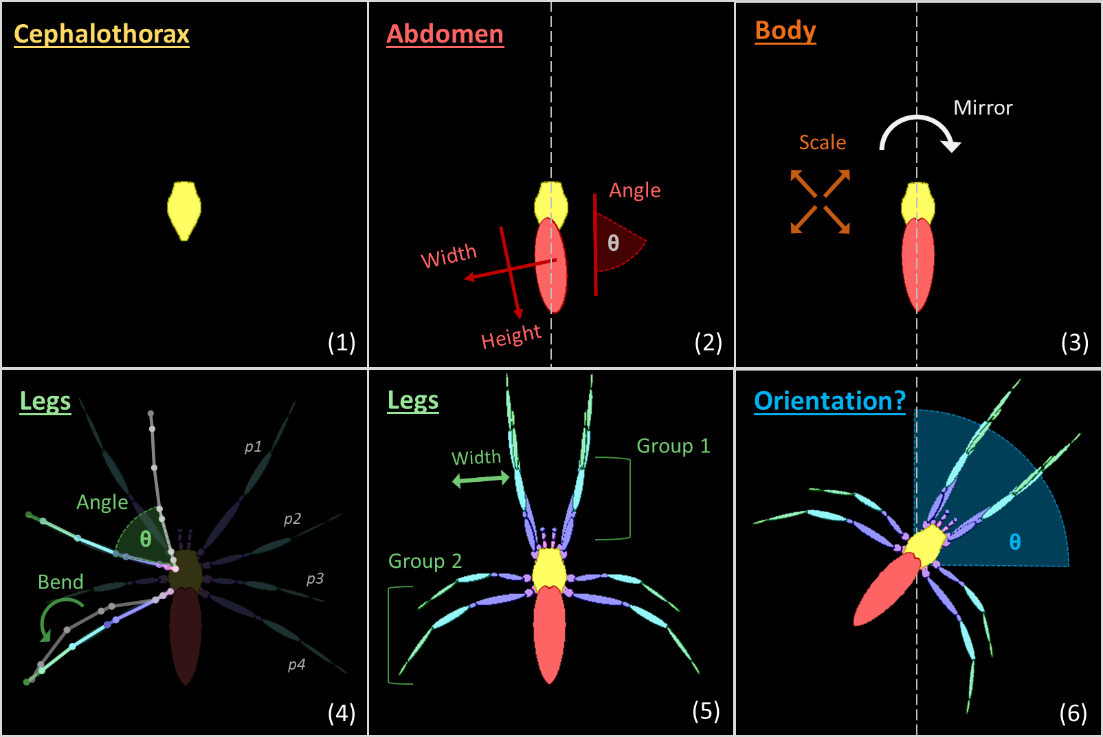

**Fig. S2. Spider mask generation.** Phases of spider mask generation. (1) the predefined cephalothorax. 2) creation of the abdomen by adjusting the width, height & angle (pointiness). (3) rescaling of the body. (4-5) generation of legs using connected splines, reangled from a base configuration. (6) Reorientation if the treatment was adaptive, note reorientation is post pattern application.

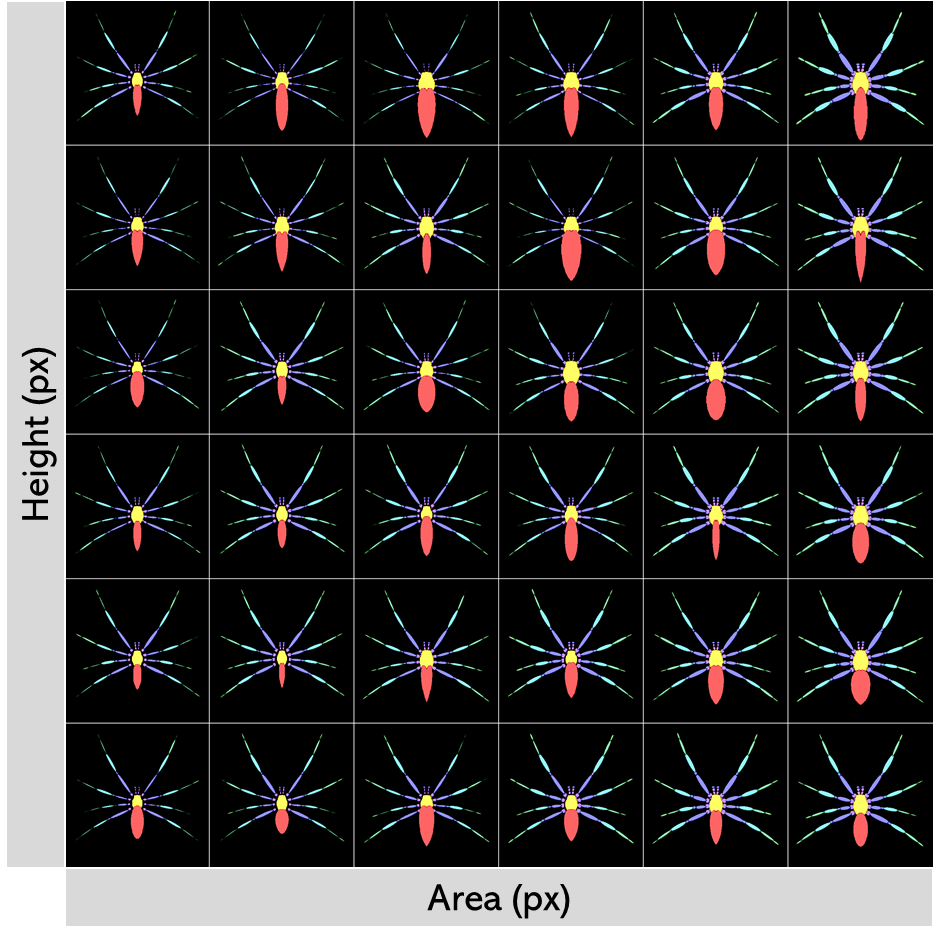

**Fig. S3. Spider scaling.** Sample spider starting population where only the abdomen shape, body width and leg width genes were allowed to vary. The spiders are distributed by the area and bounding heights of their masks, measured in pixels. In contrast, spiders with a lower area and height are smaller.

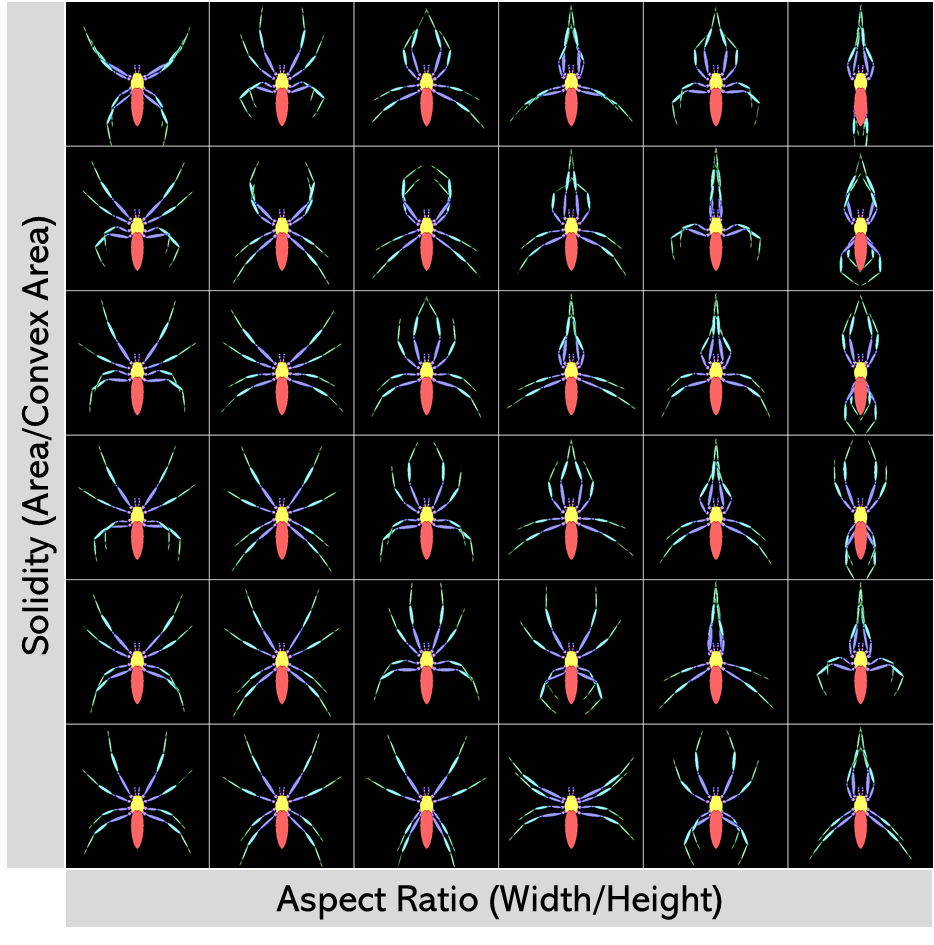

**Fig. S4. Spider postures.** Sample spider starting population where only the leg position genes were allowed to vary. The spiders are distributed by their aspect ratio and ImageJ’s measure of solidarity. Spiders with a higher solidarity and aspect ratio have a vertically stretched posture. In contrast, low solidity and aspect ratio corresponds with a circularly stretched posture.

#### SI 1.2: Spider Pattern Generation

The spiders’ colouration was generated using an experimental version of the animal pattern generator from CamoEvo. This version changed the way maculation is distributed with reaction-diffusion patterns by instead using reaction-diffusion to determine the distribution of the pattern as opposed to using a circular soft mask (Fig. S5). The new system produces complex patterns on patterns and patterns within patterns. Both the maculation and the background colours could have colour gradients instead of just the background, preventing gradients from being limited to the background. As luminance could now vary wildly across the maculation, edge enhancement was no longer determined using a ROI; instead, edge detection through gaussian blurring was used to generate the gradients. Edge enhancement could also be offset on the x and y axis to produce directionally biased edge enhancement, often seen in animals. Finally, the two speckling layers were split into before and post-edge enhancement, allowing some of the luminance speckling to have edge enhancement.

Colours were produced using a CIELAB colour space (CIE LAB; L = 0:100, A = -25:35, B=-5:50) based on the natural colour ranges of the photographed backgrounds and of spiders (Fig. S6). Each spider was comprised of four LAB colours. These colours were split into maculation and background. Colour 1 and colour 2 of each were mixed by using gaussian noise of different spatial scales as a layer mask to create internal gradients. Then the maculation and background colours were added together using the combined reaction-diffusion pattern as a layer mask, where the background used the inverse of the mask.

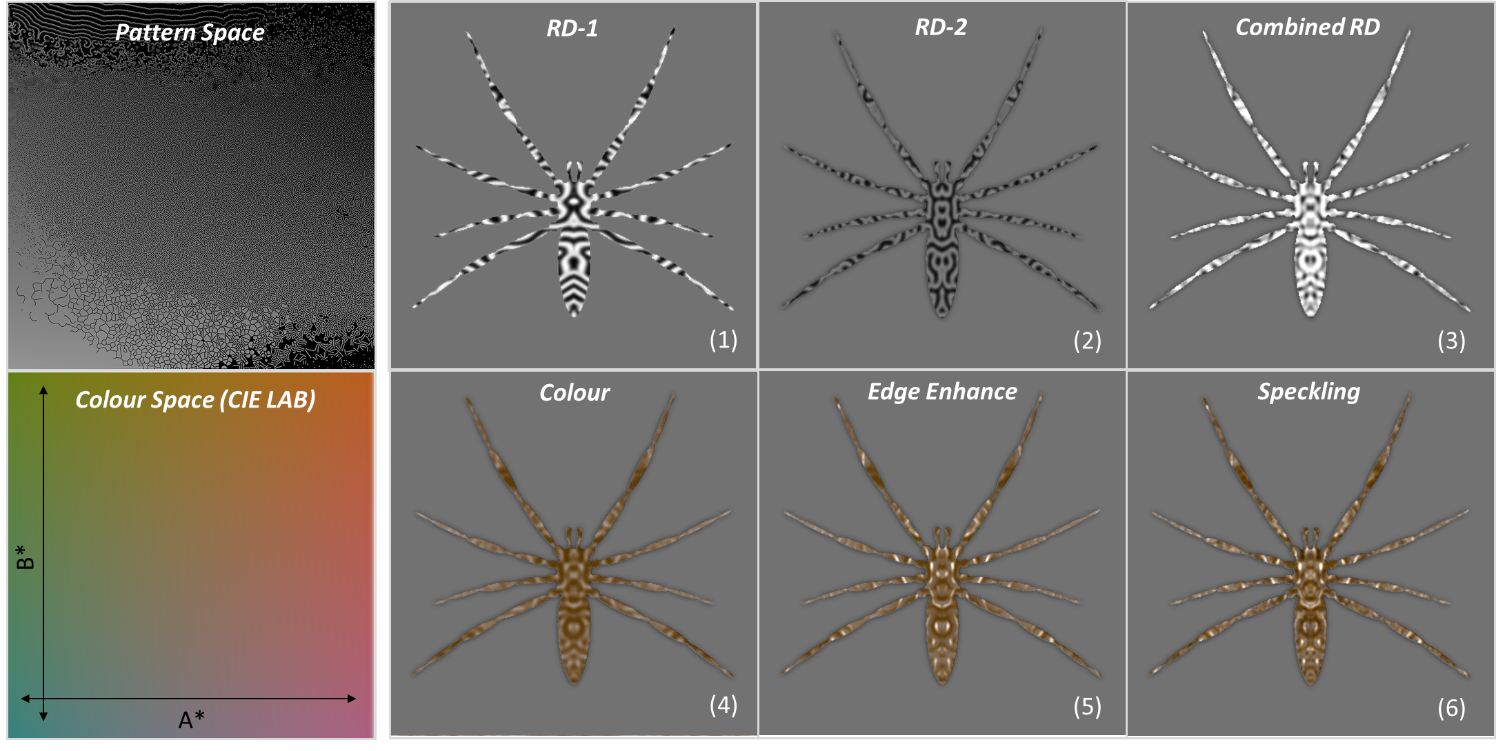

**Fig. S5. Pattern and colour generation.** Phases of spider pattern and colour generation. Spiders were created with a reaction-diffusion sheet (4000px^2^) of animal-like patterns and colours from a pre-set colour space (CIE LAB; L = 0:100, A = -25:35, B=-5:50). (1-2) sample reaction-diffusion patterns. (3) Combine the sampled reaction-diffusion. (4) Apply CIELAB colour to reaction diffusion. (5) Apply edge enhancement using luminance & saturation edge detection. (6) Add additional random speckling noise

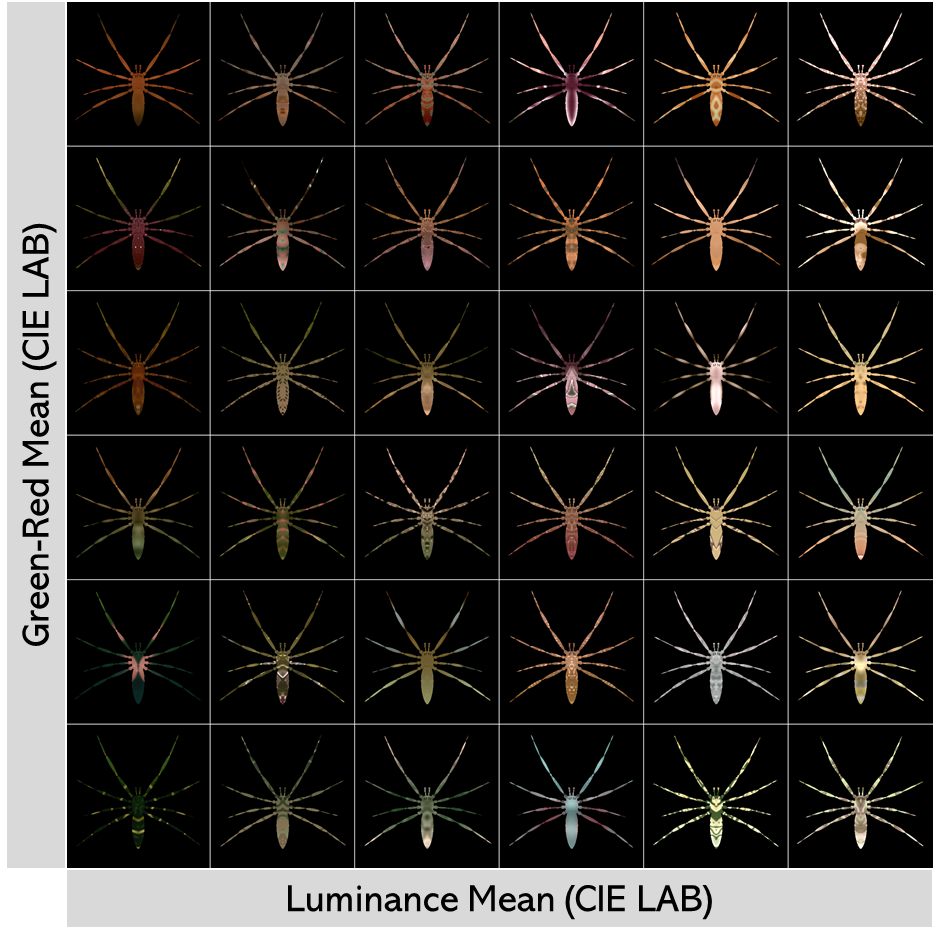

**Fig. S6. Spider patterns.** Sample spider starting population where only the pattern genes were allowed to vary. The spiders are distributed by their mean luminance (L*) and green-red colour (A*) measured in the CIELAB colour space.

#### SI 1.3: Spider Genes

As with CamoEvo, the spiders were encoded by using decimal values, ‘genes’ (see Table S1). These determined the colour, shape, pattern, posture and, in the case of the adaptive treatment, orientation of the spiders. The SpiderEvo generator uses 71 genes, divided into pattern (x20), edge enhancement (x10), colour (x12), gradient (x6), speckling (x8), leg position (x6), and body shape and orientation (x6).

**Table S1.** **Genes.** Table of all spider phenotype decimal genes.

| Section | Gene Name | Function |
| --- | --- | --- |
| PATTERN | ptn_dm1_xcp | X coordinate that pattern 1 is sampled from. |
|  | ptn_dm1_ycp | Y coordinate that pattern 1 is sampled from. |
|  | ptn_dm1_wdt | The width pattern 1 is sampled at (changes spatial scale) |
|  | ptn_dm1_asr | The aspect ratio of pattern 1 (stretches to form stripes) |
|  | ptn_dm1_agl | The rotation of pattern 1 (changes pattern angle) |
|  | ptn_dm2_xcp | X coordinate that pattern 2 is sampled from. |
|  | ptn_dm2_ycp | Y coordinate that pattern 2 is sampled from. |
|  | ptn_dm2_wdt | The width pattern 2 is sampled at (changes spatial scale) |
|  | ptn_dm2_asr | The aspect ratio of pattern 2 (stretches to form stripes) |
|  | ptn_dm2_agl | The rotation of pattern 2 (changes pattern angle) |
|  | ptn_thr_com | Changes the ratio between pattern 1-2 |
|  | ptn_thr_blr | Changes the contrast of the pattern |
|  | ptn_thr_cvr | Changes the area the pattern covers. |
|  | ptn_bil_int | Intensity of asymmetry noise applied. |
|  | ptn_bil_sig | Scale of asymmetry noise applied. |
|  | ptn_bil_ycp | Y position on asymmetry noise sheet. |
|  | ptn_si1_oft | Offset of sine wave deform of X axis (creates angle variation) |
|  | ptn_si1_sin | Intensity of sine wave deform of X axis |
|  | ptn_si2_oft | Offset of sine wave deform of Y axis (creates angle variation) |
|  | ptn_si2_sin | Intensity of sine wave deform of X axis |
| EDGE ENHANCEMENT | eeE_drk_int | L value (0 dark, 1 light) applied to negative energy regions. |
|  | eeE_drk_blr | Blur of edge enhance applied to negative energy regions. |
|  | eeE_drk_xft | X offset of edge enhance applied to negative energy regions. |
|  | eeE_drk_yft | Y offset of edge enhance applied to negative energy regions. |
|  | eeE_drk_pow | Power value applied to edge detection (changes scale) |
|  | eeE_lgt_int | L value (0 dark, 1 light) applied to positive energy regions. |
|  | eeE_lgt_blr | Blur of edge enhance applied to positive energy regions. |
|  | eeE_lgt_xft | X offset of edge enhance applied to positive energy regions. |
|  | eeE_lgt_yft | Y offset of edge enhance applied to positive energy regions. |
|  | eeE_lgt_pow | Power value applied to edge detection (changes scale) |
| COLOUR | col_mc1_lmv | Maculation colour 1, luminance channel (L*). |
|  | col_mc1_rgv | Maculation colour 1, green-red channel (a*). |
|  | col_mc1_byv | Maculation colour 1, blue-yellow channel (b*). |
|  | col_mc2_lmv | Maculation colour 2, luminance channel (L*). |
|  | col_mc2_rgv | Maculation colour 2, green-red channel (a*). |
|  | col_mc2_byv | Maculation colour 2, blue-yellow channel (b*). |
|  | col_bg1_lmv | Background colour 1, luminance channel (L*). |
|  | col_bg1_rgv | Background colour 1, green-red channel (a*). |
|  | col_bg1_byv | Background colour 1, blue-yellow channel (b*). |
|  | col_bg2_lmv | Background colour 2, luminance channel (L*). |
|  | col_bg2_rgv | Background colour 2, green-red channel (a*). |
|  | col_bg2_byv | Background colour 2, blue-yellow channel (b*). |
| COLOUR GRADIENT | grd_mac_scl | Maculation colour gradient scale. |
|  | grd_mac_sig | Maculation colour gradient blur. |
|  | grd_mac_con | Maculation colour gradient contrast. |
|  | grd_mac_ypo | Maculation colour speckle sheet y position. |
|  | grd_bkg_scl | Background colour gradient scale. |
|  | grd_bkg_sig | Background colour gradient blur. |
|  | grd_bkg_con | Background colour gradient contrast. |
|  | grd_bkg_ypo | Background colour speckle sheet y position. |
| SPECKLING | spk_st1_int | Pre edge-enhancement speckling intensity. |
|  | spk_st1_scl | Pre edge-enhancement speckling scale. |
|  | spk_st1_sig | Pre edge-enhancement speckling secondary blur. |
|  | spk_st1_con | Pre edge-enhancement speckling secondary contrast. |
|  | spk_st1_ypo | Pre edge-enhancement speckling y position. |
|  | spk_st2_int | Post edge-enhancement speckling intensity. |
|  | spk_st2_scl | Post edge-enhancement speckling scale. |
|  | spk_st2_sig | Post edge-enhancement speckling secondary blur. |
|  | spk_st2_con | Post edge-enhancement speckling secondary contrast. |
|  | spk_st2_ypo | Post edge-enhancement speckling y position. |
| LEG POSITION | leg_gen_st1 | Leg group 1 angle strength (0-100% angle dominance). |
|  | leg_gen_st2 | Leg group 2 angle strength (0-100% angle dominance). |
|  | leg_gen_ag1 | Leg group 1 intended angle. |
|  | leg_gen_ag2 | Leg group 2 intended angle. |
|  | leg_gen_bd1 | Leg group 1 bend towards centre line. |
|  | leg_gen_bd2 | Leg group 2 bend towards centre line. |
| BODY | bod_abd_wdt | Abdomen oval width. |
|  | bod_abd_hgt | Abdomen oval height. |
|  | bod_abd_agl | Abdomen oval angle (pointiness). |
|  | bod_shp_siz | Body scale. |
|  | bod_leg_wdt | Leg width/thickness. |
|  | bod_ort_agl | Body orientation (0-90 degrees). |

### SI 2: Background Image Generation

#### SI 2.0: Photography

We photographed habitats of stick spiders using an ASUS A002 smartphone and grey (4.5% and 96.2%) Spectralon reflectance standards with scale for calibration. Each background image simulates a natural branched habitat and consists of stacked layers derived from two photographs, one of sticks and the other of vegetation (Fig. S7). For this, we first collected a set of 120 sticks in woodlands located in Falmouth and Penryn (Cornwall, UK), always in sites where stick spiders (e.g., *Tetragnatha* spp.) occur. We constructed 36 different branched backgrounds with random orientation (0-360º), numbers (between 3 to 5), and types of sticks on a grey board containing the grey reflectance standards. We obtained the photographs outdoors on a sunny day by positioning the smartphone on a tripod with the lens perpendicularly to the sticks inside a Neewer diffuser tent of 1.5 x 1.5 m. We also used the smartphone on a tripod and the reflectance standards to obtain 36 blurred photos (out of focus using the manual focus function) of vegetation backgrounds in locations where stick spiders usually establish their webs. For this set of images, we standardized the position of the camera lens (parallel and 1.0-1.5 m above the ground) and the lighting (diffuse light in shaded environments). For all images, we used the manual settings of the smartphone camera, with a fixed aperture, and saved the files as Portable Network Graphics (PNG).

#### SI 2.1: Image Adjustment

Image adjustments of colour calibration, scaling, and merging layers were undertaken using the package MicaToolbox (Troscianko and Stevens 2015) of Image J, whereas removing backgrounds also required the software Adobe Photoshop (version 23.5.0, Adobe™). We generated multispectral images to calibrate the colours of vegetation and sticks in relation to the reflectance standards. Then we ran a batch scale bar calculation to determine the minimum px/mm for all the images and a macro function (Batch_Rescale.ijm) to perform the batch rescaling of all the images. We used personalized macros (Transparency.ijm and Image_Combiner.ijm) plus Adobe Photoshop to remove the background of stick images and re-cropping stick and vegetation images to 2400 px to 1800 px PNG files. We stacked a total of 36 random stick-on-vegetation pairs of images to create the branched backgrounds for the game. Stick images lacked shadows as photos were undertaken under highly diffuse light conditions.

#### SI 2.2: Spawn Map Generation

CamoEvo was choose the spider targets and backgrounds in a randomly ordered sequence and then copy them to random locations on the background images. To prevent spiders from being randomly placed beyond the sticks we created a spawn map system now incorporated into CamoEvo 2.0. Potential locations were identified by the experimenters and then using the macro (Draw_Angle.ijm), lines were drawn along the sticks and a RGB stack image was created where the R channel represented areas where the target could appear and the G channel represented the angle of the line. These could then be saved as a PNG file containing the possible coordinates and background orientations. See CamoEvo 2.0 handbook for more details: <https://github.com/GeorgeHancock471/CamoEvo-v2.0-2022_Plugins>.

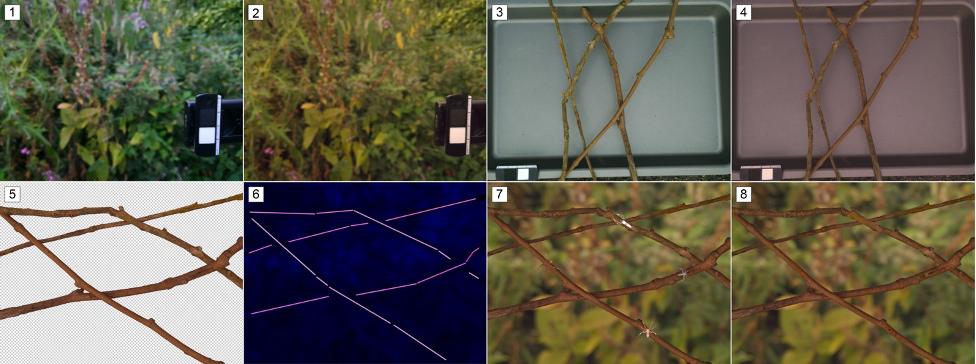

**Fig. S7. Photography and digital imaging.** Steps from image acquisition to background generation. (1) Original and (2) colour-calibrated/rescaled images of a vegetation background photographed under diffuse light in natural conditions, and (3-4) of the branches photographed under diffuse light inside an outdoor studio. (5) Transparent images of the branches. (6) Spawn maps matching the branched structure. (7-8) Example of a background (branch and vegetation), with examples of four first (7) and (8) last generation spiders, the latter occupying the same positions as the spiders of the first ones.

### SI 3: Statistics

#### SI 3.1: Effects of behaviour on camouflage

Here we provide the full list of model output for the “*Effects of behaviour on camouflage”* section of the main text (Tables S2 to S4).

**Table S2. Capture time (milliseconds) across generations**, lme4 linear mixed model output and post hoc comparisons.

| Model Output - Change in Log Capture Time (milliseconds) | | | | | | |
| --- | --- | --- | --- | --- | --- | --- |
| Predictor | Estimate | | Std.Error | DF | t Value | p Value |
| (Intercept) | 6.488 | | 0.033 | 22 | 194.57 | <0.001 |
| BehaviourGenetic | 0.010 | | 0.020 | 13990 | 0.48 | 0.629 |
| BehaviourAligned | -0.031 | | 0.020 | 13990 | -1.57 | 0.116 |
| BehaviourRandom:Generation | 0.003 | | 0.002 | 13990 | 1.64 | 0.101 |
| BehaviourGenetic:Generation | 0.028 | | 0.002 | 13990 | 14.14 | <0.001 |
| BehaviourAligned:Generation | 0.067 | | 0.002 | 13990 | 33.98 | <0.001 |
| Post Hoc Comparison | |  | | | | |
| Contrast | Estimate | | Std.Error | z ratio | p Value |  |
| Random - Genetic | -0.157 | | 0.010 | -15.08 | <0.001 |  |
| Random - Aligned | -0.351 | | 0.010 | -33.70 | <0.001 |  |
| Genetic - Aligned | -0.194 | | 0.010 | -18.62 | <0.001 |  |

**Table S3. Luminance difference across generations**, lme4 linear mixed model output and post hoc comparisons.

| Model Output - Change in Luminance Difference | | | | | |
| --- | --- | --- | --- | --- | --- |
| Predictor | Estimate | Std.Error | DF | t Value | p Value |
| (Intercept) | 17.770 | 0.393 | 72 | 45.18 | <0.001 |
| BehaviourGenetic | -1.861 | 0.400 | 13990 | -4.66 | <0.001 |
| BehaviourAligned | -1.139 | 0.400 | 13990 | -2.85 | <0.001 |
| BehaviourRandom:Generation | -0.737 | 0.040 | 13990 | -18.42 | <0.001 |
| BehaviourGenetic:Generation | -0.578 | 0.040 | 13990 | -14.45 | <0.001 |
| BehaviourAligned:Generation | -0.700 | 0.040 | 13990 | -17.51 | <0.001 |
| Post Hoc Comparison | | | | |  |
| Contrast | Estimate | Std.Error | z ratio | p Value |  |
| Random - Genetic | 0.9093 | 0.212 | 4.30 | <0.001 |  |
| Random - Aligned | 0.9199 | 0.212 | 4.35 | <0.001 |  |
| Genetic - Aligned | 0.0106 | 0.212 | 0.05 | 0.9986 |  |

**Table S4. Colour difference across generations**, lme4 linear mixed model output.

| Model Output - Change in Colour Difference | | | | | |
| --- | --- | --- | --- | --- | --- |
| Predictor | Estimate | Std.Error | DF | t Value | p Value |
| (Intercept) | 16.200 | 0.315 | 29 | 51.38 | <0.001 |
| BehaviourGenetic | -0.624 | 0.284 | 13990 | -2.20 | 0.028 |
| BehaviourAligned | -0.080 | 0.284 | 13990 | -0.28 | 0.778 |
| BehaviourRandom:Generation | -0.282 | 0.028 | 13990 | -9.92 | <0.001 |
| BehaviourGenetic:Generation | -0.259 | 0.028 | 13990 | -9.13 | <0.001 |
| BehaviourAligned:Generation | -0.385 | 0.028 | 13990 | -13.55 | <0.001 |
| Post Hoc Comparison | | | | |  |
| Contrast | Estimate | Std.Error | z ratio | p Value |  |
| Random - Genetic | 0.488 | 0.150 | 3.25 | 0.003 |  |
| Random - Aligned | 0.699 | 0.150 | 4.653 | <0.001 |  |
| Genetic - Aligned | 0.211 | 0.150 | 1.403 | 0.3393 |  |

#### SI 3.2: Effects of behaviour on posture and body shape

Here we provide the full list of model output for the “*Effects of behaviour on posture and body shape”* section of the main text (Tables S5 and S6).

**Supplemental Table 5. Spider aspect ratio across generations**, lme4 linear mixed model output and post hoc comparisons.

| Model Output - Change in Spider Aspect Ratio (Height/Width) | | | | | |
| --- | --- | --- | --- | --- | --- |
| Predictor | Estimate | Std.Error | DF | t Value | p Value |
| (Intercept) | 1.54 | 0.16 | 10.04 | 9.44 | <0.001 |
| BehaviourGenetic | -0.09 | 0.06 | 13990.00 | -1.56 | 0.12 |
| BehaviourAligned | -0.05 | 0.06 | 13990.00 | -0.95 | 0.34 |
| BehaviourRandom:Generation | 0.07 | 0.01 | 13990.00 | 12.66 | <0.001 |
| BehaviourGenetic:Generation | 0.12 | 0.01 | 13990.00 | 20.83 | <0.001 |
| BehaviourAligned:Generation | 0.29 | 0.01 | 13990.00 | 51.45 | <0.001 |
| Post Hoc Comparison | | | | |  |
| Contrast | Estimate | Std.Error | z ratio | p Value |  |
| Random - Genetic | -0.186 | 0.0293 | -6.32 | <0.001 |  |
| Random - Aligned | -1.238 | 0.0293 | -42.20 | <0.001 |  |
| Genetic - Aligned | -1.052 | 0.0293 | -35.87 | <0.001 |  |

**Table S6. Body Bounding Area across Generations**, lme4 linear mixed model output and post hoc comparisons.

| Model Output - Change in Body Bounding Area | | | | | |
| --- | --- | --- | --- | --- | --- |
| Predictor | Estimate | Std.Error | DF | t Value | p Value |
| (Intercept) | 5475.10 | 104.381 | 12 | 52.45 | <0.001 |
| BehaviourGenetic | 264.17 | 58.554 | 14025 | 4.51 | <0.001 |
| BehaviourAligned | 524.16 | 58.554 | 14025 | 8.95 | <0.001 |
| BehaviourRandom:Generation | -32.69 | 5.855 | 14025 | -5.58 | <0.001 |
| BehaviourGenetic:Generation | -75.71 | 5.855 | 14025 | -12.93 | <0.001 |
| BehaviourAligned:Generation | -35.36 | 5.855 | 14025 | -6.04 | <0.001 |
| Post Hoc Comparison | | | | |  |
| Contrast | Estimate | Std.Error | z ratio | p Value |  |
| Random - Genetic | -6.05 | 31 | -0.195 | 0.9792 |  |
| Random - Aligned | -508.1 | 31 | -16.399 | <0.001 |  |
| Genetic - Aligned | -502.04 | 31 | -16.203 | <0.001 |  |

In addition to measuring the influence of selection and behaviour on the bounding body area (cephalothorax and abdomen), we investigated how it influenced the length (bounding height) and width of the body separately. In doing so, we found that when the spiders always aligned with the background, there was no significant drop in length but there was for height (see Fig. S8 and Tables S7 and S8). Meanwhile, those where orientation was genetic evolved thinner bodies than the other treatments.

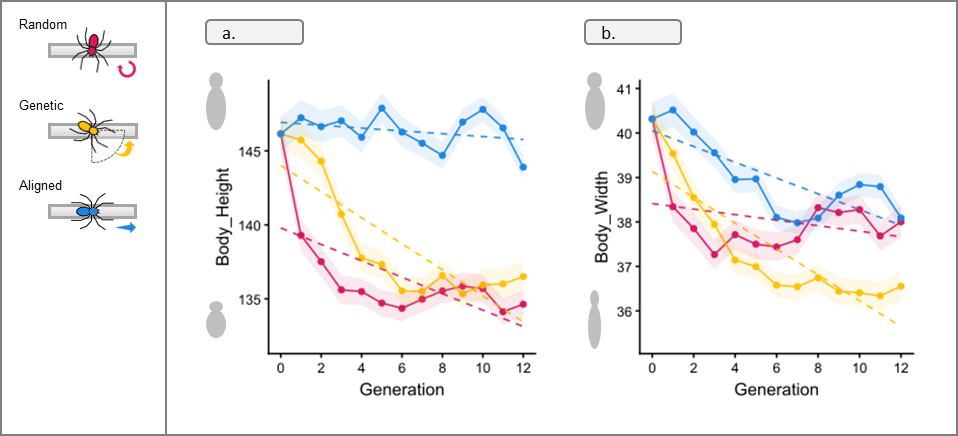

**Fig. S8.** **Change in body shape.** Difference in body shape across generations for the three behavioural treatments. (a) the change in bounding height (body length) and (b) the change in bounding width. Dashed lines show the linear trend, while the solid lines show the change in mean value for each generation and the standard error (ribbon).

**Table S7. Body bounding height across generations**, lme4 linear mixed model output and post hoc comparisons.

| Model Output - Change in Body Bounding Height | | | | | |
| --- | --- | --- | --- | --- | --- |
| Predictor | Estimate | Std.Error | DF | t Value | p Value |
| (Intercept) | 139.80 | 1.367 | 12 | 102.26 | <0.001 |
| BehaviourGenetic | 4.22 | 0.727 | 14020 | 5.80 | <0.001 |
| BehaviourAligned | 7.13 | 0.727 | 14020 | 9.81 | <0.001 |
| BehaviourRandom:Generation | -0.56 | 0.073 | 14020 | -7.65 | <0.001 |
| BehaviourGenetic:Generation | -0.88 | 0.073 | 14020 | -12.12 | <0.001 |
| BehaviourAligned:Generation | -0.10 | 0.073 | 14020 | -1.32 | 0.188 |
| Post Hoc Comparison | | | | |  |
| Contrast | Estimate | Std.Error | z ratio | p Value |  |
| Random - Genetic | -2.27 | 0.385 | -5.889 | <0.001 |  |
| Random - Aligned | -9.9 | 0.385 | -25.718 | <0.001 |  |
| Genetic - Aligned | -7.63 | 0.385 | -19.829 | <0.001 |  |

**Table S8. Body bounding width across generations**, lme4 linear mixed model output and post hoc comparisons.

| Model Output – Change in Body Bounding Width | | | | | |
| --- | --- | --- | --- | --- | --- |
| Predictor | Estimate | Std.Error | DF | t Value | p Value |
| (Intercept) | 38.41 | 0.420 | 12 | 91.40 | <0.001 |
| BehaviourGenetic | 0.72 | 0.235 | 13990 | 3.08 | <0.001 |
| BehaviourAligned | 1.64 | 0.235 | 13990 | 6.98 | <0.001 |
| BehaviourRandom:Generation | -0.06 | 0.023 | 13990 | -2.65 | 0.0081 |
| BehaviourGenetic:Generation | -0.29 | 0.023 | 13990 | -12.39 | <0.001 |
| BehaviourAligned:Generation | -0.18 | 0.023 | 13990 | -7.56 | <0.001 |
| Post Hoc Comparison | | | | |  |
| Contrast | Estimate | Std.Error | z ratio | p Value |  |
| Random - Genetic | 0.649 | 0.124 | 5.227 | <0.001 |  |
| Random - Aligned | -0.946 | 0.124 | -7.621 | <0.001 |  |
| Genetic - Aligned | -1.595 | 0.124 | -12.849 | <0.001 |  |

#### SI 3.3: Effects of angular difference on fitness

Here we provide the full list of model output for the “*Effects of angular difference on fitness”* section of the main text (Tables S9 and S10).

**Table S9. Angular difference across generations**, lme4 linear mixed model output and post hoc comparisons.

| Model Output – Change in Angular Difference | | | | | |
| --- | --- | --- | --- | --- | --- |
| Predictor | Estimate | Std.Error | DF | t Value | p Value |
| (Intercept) | 44.03 | 1.084 | 19 | 40.61 | <0.001 |
| BehaviourGenetic | -0.04 | 0.924 | 9347 | 0.04 | 0.967 |
| BehaviourRandom:Generation | 0.14 | 0.092 | 9347 | 1.52 | 0.128 |
| BehaviourGenetic:Generation | -2.58 | 0.092 | 9347 | -27.92 | <0.001 |
| Post Hoc Comparison | | | | |  |
| Contrast | Estimate | Std.Error | z ratio | p Value |  |
| Random - Genetic | 16.3 | 0.489 | 33.309 | <0.001 |  |

**Table S10. Angular difference and fitness (capture time milliseconds),** lme4 linear mixed model output and post hoc comparisons.

| Model Output - Angular Difference & Fitness | | | | | |
| --- | --- | --- | --- | --- | --- |
| Predictor | Estimate | Std.Error | DF | t Value | p Value |
| (Intercept) | -0.12 | 0.08 | 13.27 | -1.53 | 0.15 |
| Scaled AngleDifference | -0.12 | 0.01 | 9311.94 | -8.57 | <0.001 |
| BehaviourGenetic | 0.19 | 0.02 | 9309.26 | 9.31 | <0.001 |
| Scaled SpiderAspectRatio | 0.20 | 0.02 | 9323.26 | 12.44 | <0.001 |
| AngleDifference:Genetic | -0.04 | 0.02 | 9318.92 | -2.12 | 0.03 |
| AngleDifference:AspectRatio | -0.12 | 0.02 | 9315.84 | -7.95 | <0.001 |
| Genetic:AspectRatio | -0.03 | 0.02 | 9319.11 | -1.43 | 0.15 |
| AngleDifference:Genetic:AspectRatio | -0.03 | 0.02 | 9316.12 | -1.52 | 0.13 |

#### SI 3.4: Locality of background difference

To evaluate which of our background difference scales, local or immediate (Fig. S9) had the best predictive value for capture time we used AIC comparisons of models with each metric for models with either metric for CIE lightness or colour difference. Models were constructed separately for the random and aligned behavioural treatments to determine if orientation matching changed the locality of match. Models with lower AIC values were considered to be better fits to the data.

Example Model using lmer language:

lmer(log(Fitness) ~ Local_Lightness_Difference + (1|StartPop)+(1|Background)…)

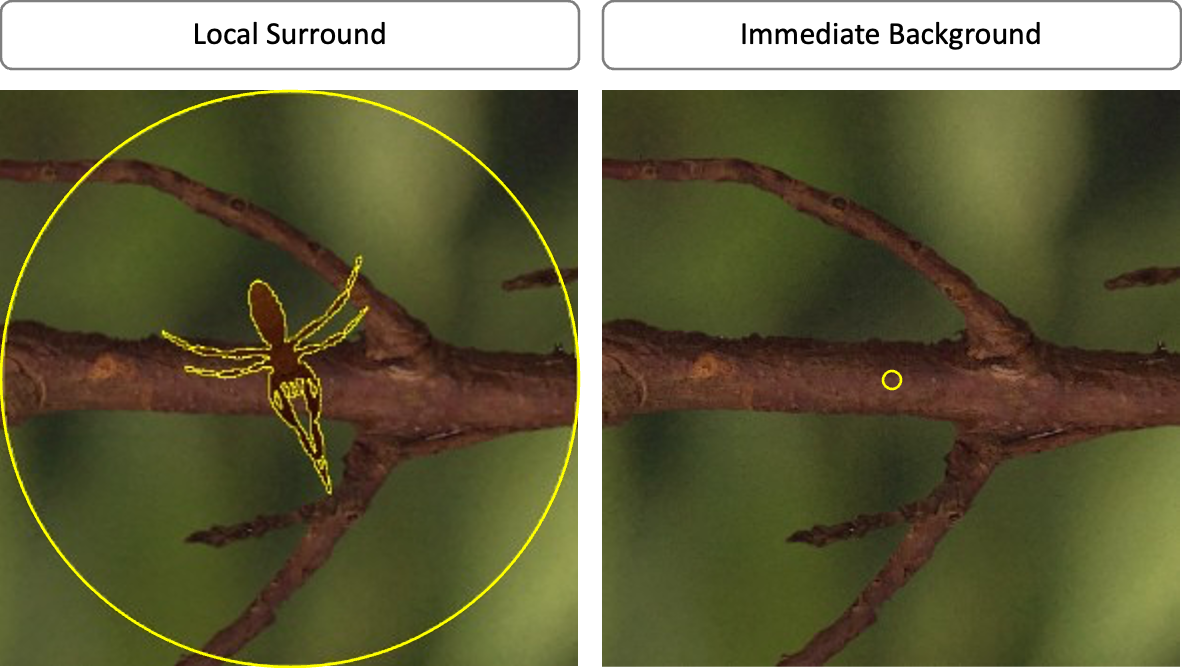

**Fig. S9. Example of background measurement selection for local surround (left) and for the immediate background (right).** Selections are shown in yellow. The local surround was an area 2x the size of the spider, excluding the spider, while the immediate background was a 10px diameter circle centred on the spider’s location.

Random Orientations:

Lightness Difference

**Local:** β = -0.006, t_4657_ = -11.00, SE = 0.001, p < 0.001**, AIC = 4706.50**

**Immediate:** β = -0.005, t _4650_ = -9.61, SE = 0.000, p < 0.001**, AIC = 4734.75**

If the behavioural treatment was random, then local luminance difference was the better predictor.

Colour Difference

**Local:** β = -0.0007, t_4657_ = -11.00, SE = 0.0008, p = 0.390**, AIC = 4824.58**

**Immediate:** β = -0.003, t _4650_ = -9.61, SE = 0.008, p < 0.001**, AIC = 4811.18**

If the behavioural treatment was random, then only the immediate background predicted fitness.

Aligned Orientations:

Lightness Difference

**Local:** β = -0.0120, t _4664_ = -12.71, SE = 0.001, p < 0.001**, AIC = 9007.01**

**Immediate:** β = -0.0146, t_4661_ = -16.53, SE = 0.000, p < 0.001**, AIC = 8900.68**

If the behavioural treatment was aligned, then immediate luminance difference was the better predictor.

Colour Difference

**Local:** β = 0.0073, t _4676_ = 5.816, SE = 0.001, p < 0.001**, AIC = 9131.38**

**Immediate:** β = -0.0191, t _4658_ = -15.17, SE = 0.001, p < 0.001**, AIC = 8940.55**

If the behavioural treatment was aligned, then local difference actually increased fitness rather than decreased fitness, while immediate difference was the better predictor.

### SI 4: Example Spiders

To make it clearer how spiders differed between behaviours we provide larger scale examples of the evolved spiders from our experiment. We show all 36 members of the populations from generations 0 and 12 for starting population 3 (Fig. S10), as well as examples for all populations at different fitness levels (Fig. S11).

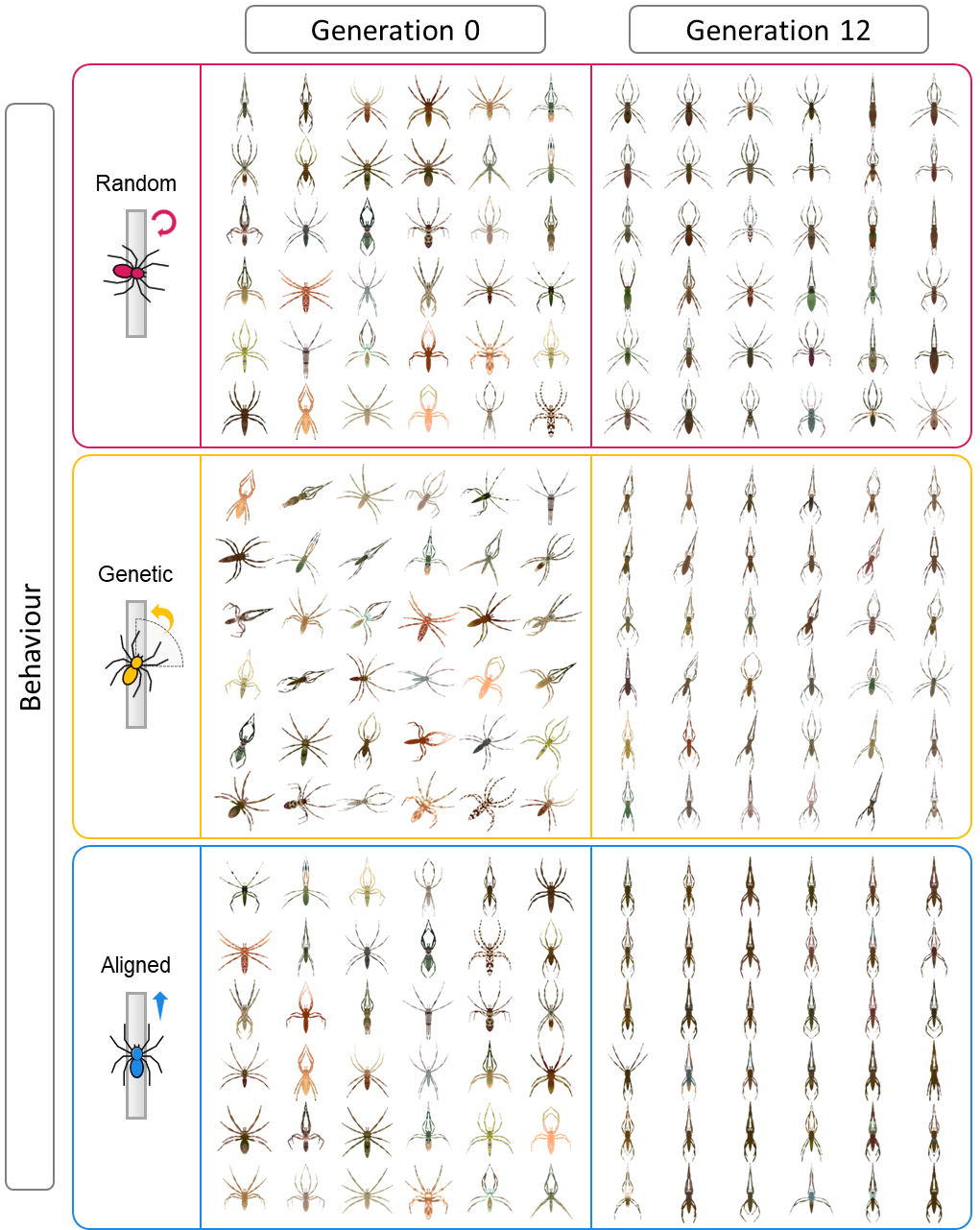

**Fig. S10. Complete populations of 36 spiders for starting population 3.** Populations are taken from the first (0) and last (12) generations and for the three different behavioural treatments. Note, only genetic is shown at different angles as the spider’s angle for the random was not heritable (instead determined by CamoEvo), while the angle for the aligned was always parallel to the substrate.

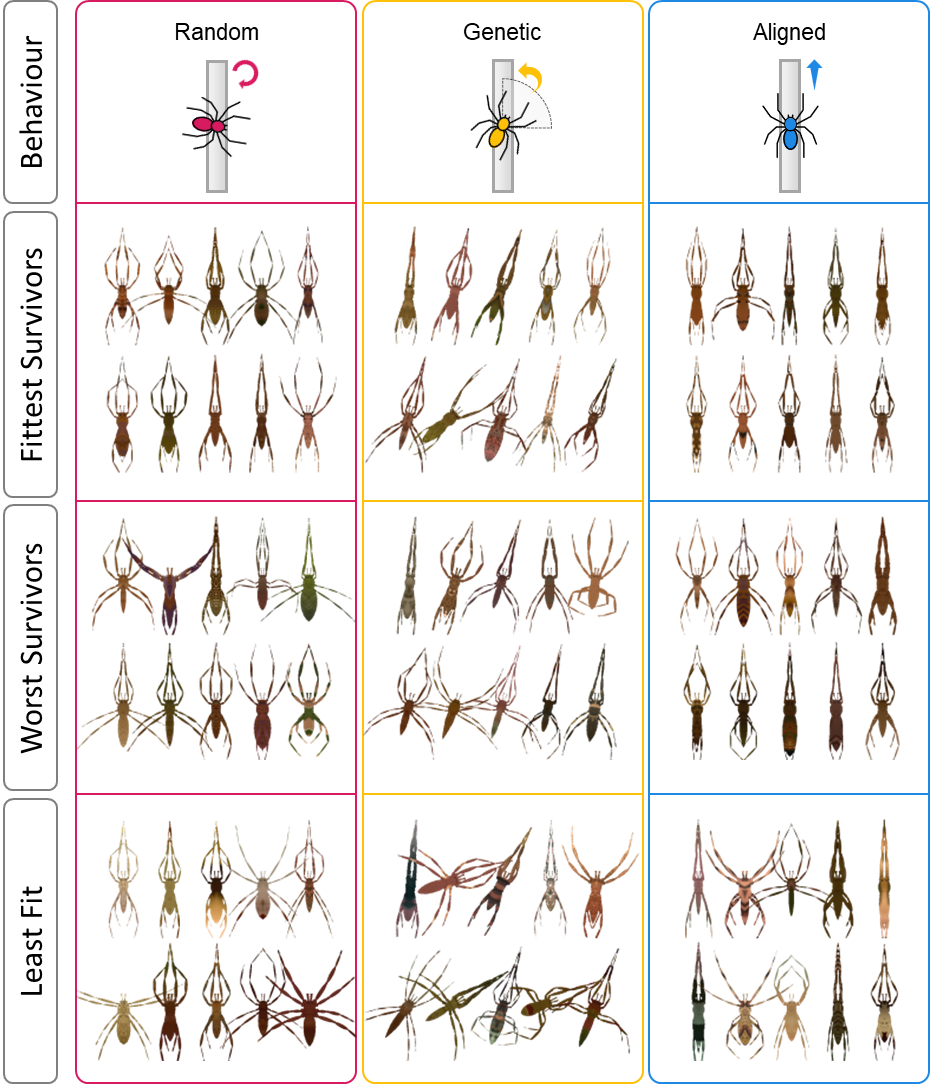

**Fig. S11. Example spiders for all 10 starting populations and behavioural treatments**. Panels include each of the fittest survivors (rank 1 for fitness), worst survivors (rank 12 for fitness), and the worst spider (rank 36 for fitness). The fitness for each spider was determined by the time taken to capture the spider. Note, only genetic is shown at different angles as the spider’s angle for the random was not heritable (instead determined by CamoEvo), while the angle for the aligned was always parallel to the substrate.

### SI 5: Example Stick Mimicry in Spiders

To characterize variation in stick-like camouflage across taxa, we compiled representative species exhibiting elongated body forms and twig-mimicking postures (Table S11). Given the high diversity of spiders and the substantial intra- and interspecific variation within the group, the table is not intended to cover all stick-like species, but rather to provide an overview of the diversity of stick-like forms across spider taxa. The total number of valid species per genus and their broad geographic distributions follow current taxonomic accounts (World Spider Catalog 2026). For each genus, we classified the representative stick-like species by (i) resting leg posture as *extended* when legs are elongated along the body axis or directed anteriorly and posteriorly, enhancing the overall linear profile, and *retracted* when legs are flexed and held close to the body, reducing appendage visibility; (ii) body orientation relative to the substrate as *aligned* when the longitudinal body axis is parallel and in direct contact with the substrate, *stick-out* when the body remains parallel but projects away from the substrate, and *detached* when the body is held away from the substrate without direct contact; and (iii) camouflage strategy as *background matching* when crypsis primarily relies on resemblance in color and pattern (including disruptive coloration) to the immediate background, and *masquerade* when individuals resemble twig elements (e.g., twig tips or broken branches). Classifications were based on scientific literature and photographic evidence, including online image databases (e.g. iNaturalist, Google Images, Instagram), using targeted keyword searches (e.g. “spider” + “twig like”, “twig mimic”, “stick-like”, “stick mimic”, “twig masquerade”, “twig camouflage”, “twig mimicry”), emphasizing the expression of the stick-like phenotype at the species level rather than generalized genus-level traits. Because several species may facultatively occupy multiple microhabitats, including branched substrates, camouflage strategies were scored exclusively from observations of individuals positioned on stick-like substrates, ensuring that classifications reflect the functional expression of crypsis in the ecological context where stick-like camouflage is most relevant. In general, extended-leg postures are consistently associated with substrate alignment and background matching, whereas retracted-leg postures are predominantly linked to masquerade and stick-out strategies.

Table S11. Diversity, distribution, and traits related to colouration (background matching, disruptive coloration and masquerade) and behaviour (body–substrate alignment and resting leg posture) of spiders living in branched environments.

| **Family** | **Genus** | **Distribution** | **Representative stick-like species** | **Resting leg posture** | **Body–substrate alignment** | **Camouflage strategy** |
| --- | --- | --- | --- | --- | --- | --- |
| Oxyopidae | *Oxyopes* | Cosmopolitan (mainly tropical–subtropical) | *O. tridens* | Extended | Aligned | Background matching |
| Philodromidae | *Tibellus* | Holarctic | *T. kibonotensis, T. maritimus, T. duttoni* | Extended | Aligned | Background matching |
| Pisauridae | *Pisaurina* | Nearctic | *P. mira* | Extended | Aligned | Background matching |
| Senoculidae | *Senoculus* | Neotropical | *S. canaliculatus, S. fimbriatus* | Extended | Aligned | Background matching |
| Theridiidae | *Brunepisinus* | Indomalayan | *B. selirong* | Extended | Aligned | Background matching |
| Thomisidae | *Monaeses* | Paleotropical | *M. israeliensis, M. paradoxus* | Extended | Aligned | Background matching |
| Thomisidae | *Borboropactus* | Paleotropical | *Borboropactus cinerascens* | Extended | Aligned | Background matching |
| Thomisidae | *Sidymella* | Australasian | *S. bicuspidata, S. longipes* | Extended | Aligned | Background matching |
| Thomisidae | *Uraarachne* | Neotropical | *Uraarachne sp.* | Extended | Aligned | Background matching |
| Araneidae | *Eustala* | Neotropical (extending into the southern Nearctic) | *E. fuscovittata, E. oblonga, E. sagana* | Retracted | Aligned | Background matching |
| Desidae | *Paramatachia* | Australasian | *Paramatachia sp.* | Retracted | Aligned | Background matching |
| Tetragnathidae | *Tetragnatha* | Cosmopolitan | *T. extensa, T. montana, T. nigrita* | Extended | Aligned | Background matching + masquerade |
| Theridiidae | *Episinus* | Cosmopolitan (mainly tropical–subtropical) | *E. angulatus, E. truncatus* | Extended | Aligned | Background matching + masquerade |
| Thomisidae | *Tmarus* | Cosmopolitan | *T. piger, T. longicaudatus, T. marmoreus* | Extended | Aligned | Background matching + masquerade |
| Deinopidae | *Deinopis* | Pantropical | *D. cylindracea, D. spinosa* | Extended | Aligned or Detached | Background matching + masquerade |
| Uloboridae | *Miagrammopes* | Pantropical (extending into subtropical regions) | *M. brevicaudus, M. orientalis, M. constrictus* | Extended | Stick-out or Detached | Masquerade |
| Theridiidae | *Ariamnes* | Pantropical (with notable island radiations) | *A. cylindrogaster, A. colubrinus, A. longissimus* | Extended | Stick-out or Detached | Masquerade |
| Theridiidae | *Rhomphaea* | Pantropical | *R. rostrata* | Retracted | Stick-out | Masquerade |
| Araneidae | *Poltys* | Paleotropical | *P. columnaris, P. laciniosus, P. noblei* | Retracted | Stick-out | Masquerade |
| Araneidae | *Dolophones* | Australasian | *D. turrigera* | Retracted | Stick-out | Masquerade |
| Araneidae | *Cyphalonotus* | Paleotropical | *C. larvatus* | Retracted | Stick-out | Masquerade |
| Araneidae | *Wixia* | Neotropical | *W. abdominalis* | Retracted | Stick-out | Masquerade |
| Araneidae | *Caerostris* | Paleotropical | *C. sexcuspidata* | Retracted | Stick-out | Masquerade |
| Archaeidae | *Eriauchenius* | Afrotropical (Madagascar) | *E. workmani* | Retracted | Stick-out | Masquerade |
